# Striated fiber assemblins build the feeding groove of an excavate flagellate

**DOI:** 10.1101/2024.03.29.587336

**Authors:** Aaron A. Heiss, Yi-Kai Fang, Anežka Konupková, Justyna Zítek, Jiří Novák, Adam Šmída, Marie Zelená, Ezra Švagr, Jiří Mikšátko, David Liebl, Priscila Peña-Diaz, Vladimír Hampl

**Affiliations:** Department of Parasitology, BIOCEV, Faculty of Science, Charles University, Vestec 252 50, Czech Republic; Kyungpook Institute of Oceanography, Kyungpook National University, Buk-gu, Daegu 41566, Republic of Korea; Laboratory of Cell Motility, Institute of Molecular Genetics of the Czech Academy of Sciences, Vídeňská 1083, Prague 142 20, Czech Republic; Faculty of Science, BIOCEV, Faculty of Science, Charles University, Vestec 252 50, Czech Republic

**Keywords:** Last Eukaryotic Common Ancestor, cytoskeleton, early eukaryotic evolution, feeding groove

## Abstract

The spatial arrangement of microtubules in the cytoskeleton in the cells of protists has been used for decades for taxonomy and phylogenetic inference at various levels. In contrast, the protein composition of non-microtubular structures is mostly unknown. Exceptions are system I fibers in algae, which are built of striated fiber assemblins (SFAs). Interestingly, SFAs are also components of a range of other, dissimilar structures, playing a role in the cortex of ciliates, cell division in apicomplexans, and adhesion of the parasite *Giardia* to the intestine. In a broad bioinformatic survey, we show the existence of three ancestral eukaryotic paralogs of SFA, and note that they are present in all “typical excavates”: small heterotrophic flagellates bearing a ventral feeding groove. In one representative, *Paratrimastix pyriformis*, we detected two SFA paralogs using specific antibodies and expansion microscopy. We show that they co-localize selectively with several microtubules and structures attached to the basal body of the posterior flagellum, namely the right microtubular root, B-fiber, C-fiber, and composite fiber. We demonstrate that one of the paralogs self-assembles in vitro into striated filaments which, under negative staining and cryo-electron microscopy, resemble system I fibers as seen in previous studies. Given the facts that all three SFA paralogs appear to be ancestral to most eukaryotic lineages, as is probably the morphology of “typical excavates” with a ventral groove, we speculate that these proteins played roles in the support and development of the feeding apparatus of the last eukaryotic common ancestor.

**SIGNIFICANCE STATEMENT:** We identified three paralogues of intermediate filament-like proteins from the family of striated fiber assemblins in the Last Eukaryotic Common Ancestor (LECA). Using expansion microscopy, we demonstrated that two of these proteins form a complex cytoskeleton that supports the feeding groove of *Paratrimastix pyriformis*. It is widely accepted that this flagellated protist retains the morphological and feeding characteristics of ancestral eukaryotes. Therefore, our results suggest that the striated fiber assemblin proteins, which form diverse structures in various extant eukaryotes, were initially components of the feeding apparatus in LECA.

## INTRODUCTION

The eukaryotic cytoskeleton is a complex system comprising microtubules (MTs), microfilaments, and various fibrous structures^1^. In the context of unicellular eukaryotes (protists), it can be the primary determinant of the morphology of an entire organism. It is sufficiently conserved as to have long been recognized as taxonomically significant at deep phylogenetic levels^2,3,4,5^. In most protists, the core of the cytoskeleton is the flagellar apparatus, comprising the basal bodies and the microtubular and fibrous structures directly associated with them. Some of these structures, termed flagellar roots, can extend through the full length of the cell, often providing an armature stabilizing the cell’s shape^1^.

The microtubular components of the cytoskeleton have received the bulk of attention^4,6^. Microtubular roots are suspected to be homologous in different lineages if they appear in a similar arrangement both to the basal bodies and to each other, and extend along similar paths through the cell^4^. Associations with non-microtubular fibrous structures are also significant^3,5^. If these fibrous structures are composed of similar materials, the likelihood of homology between the roots increases considerably^3^.

System I fibers^1,7^ are common cytoskeletal structures associated with microtubular roots in green algae, built of the protein striated fiber assemblin (SFA) protein^8^. SFA genes are much more widespread amongst eukaryotes^9^ but have been characterized only in a few systems, namely, as the components of the microribbons of the adhesive disc in *Giardia*^10^, as kinetodesmal fibers in ciliates^9,11^, as fibers associated with basal bodies and microtubules in oomycetes^12^, and with the cell division apparatus in apicomplexans^13,14^.

In this study, we expand the taxonomic range of eukaryotes with SFA genes and localize them in the cell of *Paratrimastix pyriformis*. This organism bears a body plan resembling one of the probable common ancestor of most or all eukaryote lineages^4,15,16,17,18^. A prominent structure of this body plan is a ventral feeding groove, supported by three microtubular roots attached to the posterior basal body, and associated with several fibers of unknown, but probably proteinaceous, composition. This “typical excavate” morphology is present in many extant members of three distantly related lineages, Discoba, Malawimonadida, and Metamonada (the latter including *P. pyriformis*). The small size of “typical excavate” cells (≤20 µm long) prevents clear localization of small-scale structures such as microtubules under conventional fluorescence microscopy. Thus, we used expansion microscopy, determining that three non-microtubular fibers in the feeding groove of *P. pyriformis* are composed of SFA, and also that SFA specifically associates with some microtubules of the groove. We also confirm, using transmission electron microscopy, that the fibers that SFA constructs form spontaneously in vitro, and have banding patterns consistent with those of system I fibers and their homologs in other organisms. This suggests that the original function of SFA may be connected to the excavate feeding groove and opens the way to confirm homologies between groove components in phylogenetically distant organisms.

## RESULTS

### Phylogenetic Analysis

We conducted a phylogenetic analysis of 302 striated fiber assemblin (SFA) homologs from 128 taxa, which separated the proteins into two major clades (Figure S1). One of these clades, corresponding to ‘Group 1’ in Soh et al.^9^, contained (a) all members of the green algal lineage included in the analysis; (b) all of the three previously studied SFA homologs in *Giardia*; and (c) all of the previously identified apicomplexan and stramenopile SFA homologs. The other clade (corresponding to ‘Group 2’ in Soh et al.^9^) contained almost all of the ciliate sequences and about half of the fornicate sequences; other ‘kingdom-scale’ lineages were far less well-represented. Notably absent from either group were lineages comprising the ‘megagroup’ Amorphea^19^.

To improve the phylogenetic signal within groups, we analyzed each separately. With only three exceptions, all of the lineages present in Group 1 were monophyletic and received significant statistical support (Figure 1). The exceptions were (i) cryptists, of which *Palpitomonas* branched separately from the remainder, (ii) ciliates, which branched separately from all other members of the SAR clade, and (iii) Metamonada, of which Fornicata and Preaxostyla branched separately in all analyses. The backbone of the Group 1 tree was unsupported. Four lineages showed internal paralogy. Haptophytes and alveolates (without ciliates) each independently evidenced a clear duplication of SFA into two highly supported subclades, each with comparable taxon composition. Diplomonads, retortamonads, and *Dysnectes* (within fornicates) formed three subclades, corresponding to all three of the SFA paralogs previously studied in *Giardia* (β-giardin, δ-giardin, and SALP-1); these subclades received varying levels of ML support, but maximal support from Bayesian analysis. Finally, we found two copies of Group 1 SFA in most species of Preaxostyla.

**Figure 1:**
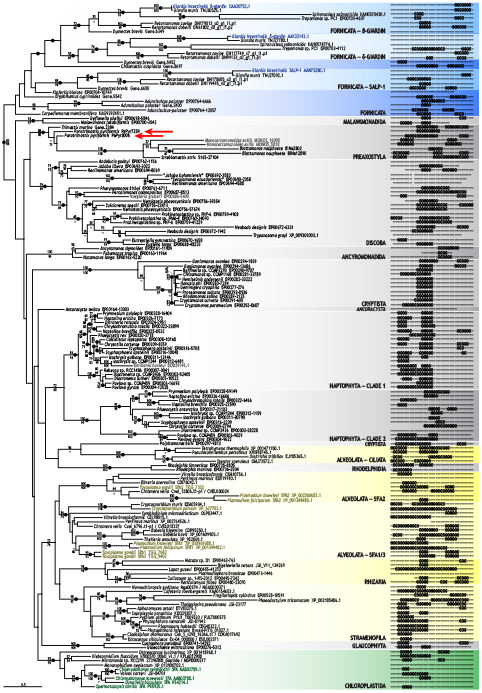
Phylogenetic tree of Group 1 paralogs of striated fiber assemblin. Tree was generated from maximum-likelihood search using GTR+Γ model as implemented in RAxML; topology is from the best of 100 starting random-taxon-addition parsimony trees. Support values along branches are from 1000 nonparametric bootstrap replicates (above branches) and from 200 bootstrap replicates using the LG+PMSF+C20+G model with 100 starting trees per replicate (below). Values < 50% are indicated by a dash (‘–’), or omitted when both analyses returned < 50%. Black circles in branches indicate full support from Bayesian posterior probability; unlabelled branches were not supported by Bayesian analysis (probability < 0.95). Colored background highlights previously characterized lineages; colored sequence names indicate previously investigated proteins. Red arrows indicate homologs used in present study. Figures at right are mappings of predicted coiled-coil domains onto sequence, with each character indicating seven amino acid sites (the number of sites generating a single coil); ‘@’ indicates location of probable coiled-coil domains, ‘-’ indicates sequence with no coiled-coil domain prediction; absence of symbol within mapping indicates gap in alignment (i.e., deletion); absence of symbol at ends of mapping indicates sequence not available. See Figures S1 and S2 for more information.

The Group 2 tree showed two clear subclades termed ‘Group 2a’ and ‘Group 2b’ according to Soh et al.^9^ The backbones of these subclades were unsupported (Figure S2), and support for individual ‘kingdom-scale’ lineages varied. The composition of each of the Group 2 clades was dominated by fornicates and ciliates, with extensive paralogy within the ciliates. The majority of the sequences from fornicates were to be found in Group 2a, with *G. intestinalis* appearing in both groups. Most other lineages also had paralogs within Group 2, although in lower numbers overall. The most prevalent of these were non-ciliate alveolates, discobids, and preaxostylans.

### Structural Analysis

Secondary-structure predictions in SFA proteins identified only two consistently appearing domain types in SFA proteins: coiled-coil motifs and low-complexity regions. The latter generally coincided with the former, had lower confidence scores, and were not given further consideration. The former is thought to be characteristic of SFA proteins^20^, but despite this, consistent positional patterns of predicted coiled-coil domains were visible neither on the large scale nor within the three major clades (Figures 1 and S2). However, patterns did appear in some individual lineages, most prominently in cryptists and haptophytes (Figure 1).

We prepared recombinant forms of two SFA homologs of *Paratrimastix pyriformis*: PaPyr7259 and PaPyr8006 (P7 and P8, respectively), from Group 1 (Figure 1, arrows). Both exhibited some degree of self-assembly *in vitro*, which manifested as gelation in the case of P8 and as a turbid solution containing small clumps in the case of P7. To assess the nature and ultrastructural features of P7 and P8 assemblies *in vitro*, we processed both samples for analysis by transmission electron microscopy, first by fresh renaturation of urea-treated proteins and then by negative staining or cryo-EM. Negative staining showed that the P7 protein assembled into distinctively short linear structures (presumably oligomers) with a uniform length of 30-40 nm and discernible globular density bulging at one or both ends of the structure (Figure 2A). These P7 assemblies were closely reminiscent of elementary subunits of intermediate filaments, such as those reported for lamin B2^21^. In contrast, P8 assembled into substantially long rope-like filament bundles, often entangled into clusters (Figure 2B) with a characteristic (predominant) bundle width of 11.5 ± 1.0 nm, reminiscent of the supercoiled sheet structure of intermediate filaments (typical width of 8–12 nm). A substantial subpopulation of filament bundles exhibited characteristic striations: a repetitive ultrastructural feature with 34.6 ± 2.2 nm periodicity along the longitudinal axis. These were composed of stretches of traceable linear filaments (∼20 nm long) interspaced with a segment of (likely globular) non-linear protein domains (∼15 nm long), wherein the latter were markedly wider (14.0 ± 1.4 nm). Similar features (a pronounced axial beading repeat) were previously observed in lamin A filaments and in paracrystalline lamin A fibers^22^. Interestingly, even-thicker bundles and laterally aligned sheets of protofilaments were found in a sample composed of a mixture of P7 and P8 (Figure 2C–D), which we analyzed to assess the potential co-assembly of P7 and P8.

**Figure 2.**
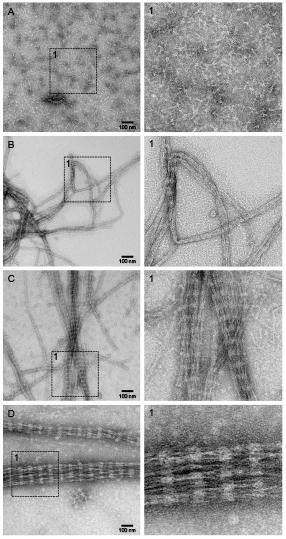
Ultrastructural analysis of P7, P8, and P7+P8 protein samples by negative staining and transmission electron microscopy. **A:** Homogeneous distribution of P7 (likely dimers or oligomers) with globular (head) and stretched (tail) domains discernible in lateral projections after adhesion on grid surface. **B:** Protofilaments and bundles assembled from P8. **C, D:** Mixture of P7 and P8 assembled into thick, laterally-aligned bundles or two-dimensional sheets with regular striation pattern. All images are pairs of overview images (on left) with enlarged details (on right). Scale bar = 100 µm. See Figure S3 for more information.

We also employed cryo-electron microscopy to analyze the structure of P7 and P8 in their intact, fully hydrated, and close-to-native form, while suspended in vitrified ice. This approach should minimize the potential for (i) distortion artifacts induced by the adhesion of protein assemblies to the solid support of the grid, (ii) artifacts related to negative staining by heavy metals, and (iii) surface tension-related drying artifacts. The results (Figure S3) confirmed the majority of findings from the negative staining and provided some additional insight. The P7 oligomers appeared less regular on cryo-EM images, likely due to the preservation of their flexible architecture and random spatial orientation within the 3D volume of the vitrified sample, seen as multiple projection angles (Fig 3A). Cryo-EM of the P8 sample revealed similar features as observed by negative staining, including the formation of protofilament bundles (Figure S3B). However, we found that the co-assembly of P7 and P8 resulted not only in the formation of planar sheets from laterally aligned bundles but also in the formation of a 3D structure composed of equilateral triangles (Figures S3C,D), a structure that may not have been preserved or discernible in our negative-stained samples.

**Figure 3:**
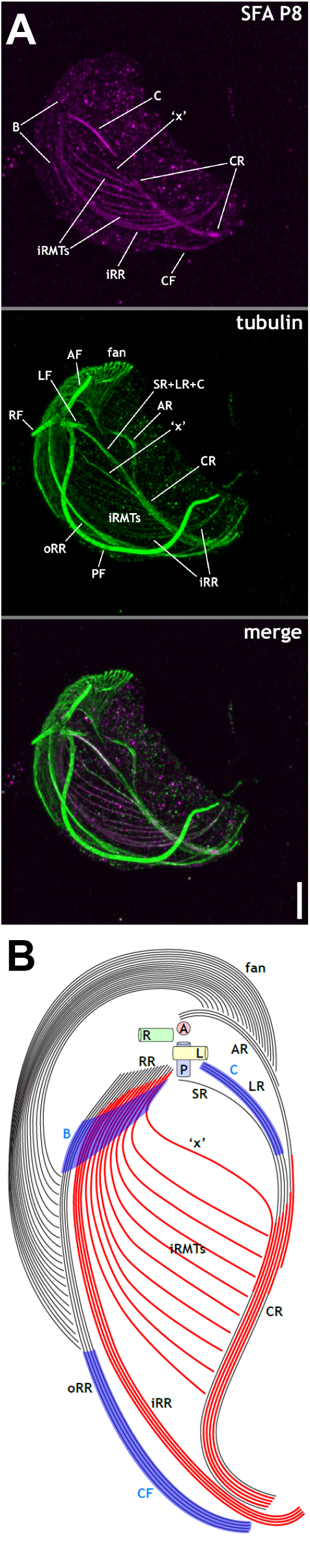
Expansion microcopy of whole cell of *Paratrimastix pyriformis*. **A:** Cell viewed from ventral-right side stained with antibodies to structural proteins; magenta **-** striated fiber assemblin paralog *PaPyr8006*; green - tubulin. Note that all flagella except the posterior flagellum have broken off close to their insertion points. Note also that singlet root is not visible as independent structure. **B:** Summary diagram of microtubular and striated-fiber-assemblin-based structures in *Paratrimastix pyriformis*. Cell is oriented with anterior to top of page and cell’s left to viewer’s right. Cylinders at cell’s anterior represent basal bodies. Plain black lines indicate microtubules (MTs) without associated striated fiber assemblin (SFA) protein; thicker red lines indicate MTs with associated SFA; blue regions indicate SFA-based structures extending beyond individual MTs. Key to annotations: ‘AF’ = anterior flagellum; ‘AR’ = anterior root; ‘B’ = B-fiber; ‘C’ = C-fiber; ‘CF’ = composite fiber; ‘CR’ = compound root; ‘fan’ = dorsal fan; ‘iRMT’ = inner right root microtubules; ‘iRR’ = inner right root; ‘LF’ = left flagellum; ‘PF’ = posterior flagellum; ‘oRR’ = outer right root; ‘RF’ = right flagellum; ‘SR+LR+C’ = compound root comprising singlet root, left root and C fibre; ‘x’ = ‘x’ microtubule in iRMT. Scale bar ∼ 2 µm (approximated based upon the diameter of flagellar axonemes and size of cell). See Figures S4 and S5 for more information.

### Localization in the cell

We raised polyclonal antibodies against recombinant P7 and P8 and confirmed that they specifically recognized bands of expected sizes in western blots of whole cell lysates (Figure S4). Co-immunoprecipitation showed that each antibody precipitated both paralogs (Tables S1, S2), even in the presence of RIPA buffer, indicating that the co-immunoprecipitation was caused by a cross-reaction of the anti-P7 antibody to the P8 protein and vice versa (Table S3). Co-immunoprecipitation also detected several common contaminants, such as ribosomal proteins, but consistently detected alpha-tubulin and, in the case of P8, beta-tubulin.

Both anti-P7 and anti-P8 antibodies produced specific immunofluorescent signals in expanded cells of *P. pyriformis* that were also immunolabelled with anti-alpha-tubulin antibody (Figures 3A, 4, S5; Videos S1-S3; summary diagram in Figure 3B). The anti-tubulin antibody labeled flagellar axonemes and basal bodies (BBs) prominently, and the preparations were sufficient to distinguish individual MTs’ paths within cells, albeit not always to quantify numbers of MTs when they were close together. Cells were preserved with varying degrees of distortion. In general, the better-preserved cells were less well-stained, and flagella were frequently broken off proximal to the cell (e.g., Figure 3A). Microtubules within cells could usually be identified with previously described structures, especially at the anterior part of the cell, in which their arrangements were consistent between cells. However, the posterior arrangement of MTs, while consistent in some cells, was highly variable in others, often appearing chaotic. Our description is therefore based on the positions, orientations, and connections most consistently seen in multiple less-disrupted cells.

Near the BBs, the ventrally projecting anterior flagellum was associated with an anterior microtubular root (AR), which arced over the anterior side of its BB in a broad leftward and posterior curve (Figures 3A, 4A, 4B, S5). The AR nucleated a conspicuous dorsal fan comprising a radiating array of ∼20 microtubules (MTs) (Figure 4A, B), although at least one cell had more (∼35 in Figure S4). Most of these were evenly spaced, with occasional and inconsistently placed gaps. The dorsal fan followed the outline of the anterior part of the cell, initially projecting dorsally and slightly posteriorly (Figure 3A, S5), then curving posteriorly and leftward (Figures 3A, 4B, 4C, S5), and finally posteriorly and rightward (Figures 4D, E), abutting and terminating at the outer edge of the posterior right root (see below), about halfway down the cell’s length (Figures 4D, E). The fan thus presented as a ribbon-like helix, curving ∼¾ around the cell from its origin (Figure 3B).

**Figure 4:**
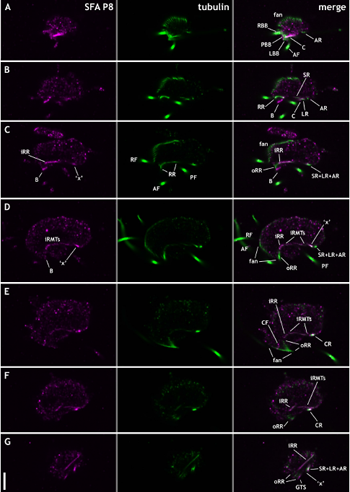
Optical slices from expansion microscopy of cell of *Paratrimastix pyriformis*. Cell is viewed in cross-section, with dorsal to top of page and cell’s left to viewer’s right. Slices are spaced ∼2–4 µm apart. **Left panels:** striated fiber assemblin paralog *PaPyr8006*; **middle panels:** tubulin; **right panels:** merged images. Key to annotations: ‘ABB’ = anterior basal body; ‘AF’ = anterior flagellum; ‘AR’ = anterior root; ‘B’ = B-fiber; ‘C’ = C-fiber; ‘CF’ = composite fiber; ‘CR’ = compound root; ‘fan’ = dorsal fan; ‘iRMTs’ = inner right-root microtubules; ‘iRR’ = inner right root; ‘LBB’ = left basal body; ‘LF’ = left flagellum; ‘LR’ = left root; ‘PBB’ = posterior basal body; ‘PF’ = posterior flagellum; ‘oRR’ = outer right root; ‘RBB’ = right basal body; ‘RF’ = right flagellum; ‘RR’ = right root; ‘SR’ = singlet root; ‘SR+LR+C’ = compound root comprising singlet root, left root and C fibre; ‘x’ = ‘x’ microtubule in iRMT. Arrow indicates initial split between iRR and iRMTs. Scale bar ∼ 2 µm (approximated based upon diameter of flagellar axonemes and size of cell). See Video S3 for more information.

The left and right posterior roots (‘LR’ and ‘RR’, respectively) each initially comprised a small but indeterminate number of tightly spaced parallel MTs (Figure 4B). A single MT, the singlet root (SR), originated near the RR, angled to the left, and joined the LR (Figure 4B, 4C; Video S3). This junction was located close to the SR’s point of origin, such that in some cells the SR was difficult to perceive as an independent entity at all (e.g., Figure 3A; Video S3). Posterior to this point, the LR and SR ran along the left ventral margin of the cell, being joined at about one-third of the length of the cell by the AR (Figure 3A; 4B, 4C; S5; Videos S1-S3).

The RR widened adjacent to its point of origin by acquiring several additional MTs along its outer (right) side. Not very far distal (posterior) to this, it split into two distinct but closely apposed subunits, the inner and outer RR (iRR and oRR, respectively), the oRR comprising notably fewer (∼4) MTs (Figure 4C; Video S3). While the MTs of the oRR remained close together, those of the iRR were spaced one to two MT diameters apart, not much less than the space comprising the split between the iRR and oRR. This often made the split difficult to discern from the tubulin stain alone. Individual MTs frayed off the iRR to the left at more or less regular intervals, forming a widely spaced set of inner RR MTs (iRMTs). These spanned the distance diagonally between “unfrayed” MTs in the iRR and the LR and its associated structures (Figure 3A, S5). The first MT to split off from the inner part of the iRR, here termed “x”, extended leftward and curved to join the LR, SR, and AR in parallel (Figure 3A, S5, 4D); posterior to that point, these MTs appeared as a single structure, which we termed the ‘compound root’ (‘CR’) (Figure 3A, S5, 4E, 4F). Other iRMTs (∼9 in total) either abutted and terminated at the CR or, more posteriorly, passed close by the CR to terminate independently not far beyond it (Figure 3A, S5). At the posterior end of the cell, the remaining ∼4 MTs of the iRR remained spaced roughly one MT width apart from each other curving leftward and then anteriorly for a small distance (Figures 3, 4F, 4G). The oRR, in contrast, remained either parallel to the CR or at a close angle to it (Figures 3, 4).

Consistent with our co-IP results, P7 and P8 antibodies showed similar localization patterns; the one from P8 is shown in Figures 3A and 4, Videos S1, S3, while the one from P7 is shown in Figure S5 and Video S2. One of the most conspicuous features was the colocalization of P7 and P8 with the MTs of the iRR along their whole lengths (Figure 3, S5). While the LR and SR were unlabeled along their anterior length, they acquired an SFA signal immediately anterior to their being joined by “x”. Posterior to that point, the SFA signal appeared only on the LR+SR MTs, and, when MT identity could be determined, not on “x” (Figure 3A; 4G).

Non-tubulin-based structures were also labeled by our SFA antibodies. One very strong signal, was to an anteroposteriorly oriented root-like structure originating near the posterior basal body (PBB). This was 2.5–3.0 µm long, and often sufficiently close to the LR as to appear part of it. This could correspond to the C-fiber as described by O’Kelly et al.^23^. In many cells, however, this structure diverged ventrally from the LR at a narrow angle (Figure 3A; 4A, 4B; Videos S1, S2), unlike the C-fiber as described previously in electron microscopy^23^.

Another structure (300–500 nm in width) was detected at the anterior of the cell, adjacent to but separate from any MTs (Figure 3B, 4B, 4C). This structure was less apparent in cells with their anteroposterior axes close to the plane of the optical section but could be made out distinctly in cells oriented perpendicularly to the plane of the section (as in Figure 4). It described a sheet running from the junction of the RR and the PBB diagonally posteriorly and rightward, to terminate about a third of the way through the cell at the oRR. Its posterior point of termination against the oRR was just anterior to or coincident with the abutment and termination of the anteriormost MT of the dorsal fan (Figure 3B). This sheet-like structure corresponds positionally to the B-fiber described by O’Kelly et al.^23^.

Finally, a structure tagged by SFA but not by tubulin antibodies appeared at the posterior of the cell, on the outside face of the oRR. This was a ribbonlike structure that originated just anterior to the closing of the groove (Figure 4E), and about at the same point as the posteriormost MT of the fan terminating against the oRR (Figure 3B, 4E). It ran through the end of the oRR (Figure 3B, 4G).

The bulk of this structure is in the same position, and is of a comparable shape, to the composite fiber described in *P. eleionoma* by Simpson et al.^22^.

## DISCUSSION

Our work builds on the spectrum of lineages shown in previous studies to have SFA homologs (most recently Nabi et al.^11^ and Soh et al.^9^). We confirm the observation of Soh et al.^9^ that eukaryotic SFAs form three major clades (Group 1, Group 2a, and Group 2b), and based on their distribution, we infer that all three were present in their common ancestor (Figure S6). Interestingly, at the deepest scale of eukaryotic diversity, there is (yet) no evidence of SFAs in any members of Amorphea (the part of the tree of eukaryotes containing the ‘supergroups’ Amoebozoa and Opisthokonta, the latter including fungi and animals). All three clades of SFAs appear in Discoba, Metamonada, and Malawimonadida (collectively, the “excavates”). Each of these lineages, or a combination of them, has generally been supposed to be closer to the eukaryotic root than Amorphea and Diaphoretickes^15,16,18,25^ (but note Cerón-Romero et al.^17^).

Notably, we found no SFA homologs in Hemimastigophora, which branches deeply within Diaphoretickes^26,27^. Hemimastigophorans have intriguingly ciliate-like cellular architectures^28^. Their lack of SFAs deepens the intrigue, given that ciliates have a staggering amount of within- and between-lineage paralogy of SFA genes. This implies that the similarity in cytoskeletal architecture between these two phylogenetically distant lineages is truly convergent. It should be interesting to find out what proteins constitute the fibers in hemimastigophorans that are analogous to those in ciliates.

The ultrastructure of *P. pyriformis* was previously studied using serial-section transmission electron microscopy (TEM) by Brugerolle and Patterson^29^ (as “*Trimastix convexa*) and by O’Kelly et al.^23^. Importantly, one of the strains studied by O’Kelly et al.^23^ (“RCP” = ATCC 50562) is the same as the one used in our work. One of the primary characteristics of “typical excavate” architecture is a posterior right root (RR or R2) that splits prominently, close to its point of origin, into two comparably sized subroots^3^. This contrasts with the split in *P. pyriformis*’s RR, however, which is offset and never more than a few MT widths wide (Figures 4D, E). The split is sufficiently subtle that it was unobserved by Brugerolle and Patterson^29^, who interpreted the RR as a single unit from which iRMTs frayed directly, an arrangement illustrated by O’Kelly et al.^23^. Our ExM results confirm this description but also indicate that not all of the iRR MTs fray (Figures 3A, S5, 4C–G; Videos S1–S3).

The posterior end of the *P. pyriformis* cell received little attention in either previous account. Brugerolle and Patterson^29^ showed several TEMs longitudinally or obliquely oriented through the cell that included the feeding groove closing at the cell’s posterior but offered no written description.

O’Kelly et al.^23^ only published TEMs of the anterior-to-middle part of the cell, although they did state that the LR and oRR “reunited at the posterior end of the ventral groove” (p. 155). Our work suggests that the reunion between LR and oRR is more complex than that, with three converging groups of MTs: the CR (itself an assembly of MTs from different sources), iRR, and oRR. Furthermore, although the oRR and CR do closely follow the same path at the posterior of the cell, they do not merge, while the iRR — positioned between them for most of their length — deviates significantly from that path (Figures 3B, 4G).

Another consistent observation in our samples was that the RR comprised no more than 20 MTs in total: 14–15 iRR MTs (including the “x” MT and ∼9 iRMTs) and 4–5 oRR MTs. This was counted in seven cells, and proportionally consistent with >40 others. In contrast, both Brugerolle and Patterson^29^ and O’Kelly et al.^23^ describe ∼30 MTs in the whole RR, although, oddly, their micrographs show ∼35 and 40, respectively. As noted, O’Kelly et al.^23^ studied the same strain as we did. The possibility of our preparation eliminating ¼ to ½ of the MTs in the RR is undermined by the consistent size of the RR that we found among our cells. In addition, the consistent position of the outer edge of the B-fiber and the abutment of the dorsal fan led us to suspect that no MTs were lost in the course of our preparation. Observation under TEM should confirm or refute this interpretation.

Strictly speaking, exact colocalization between MTs and SFA-based structures was only found in the RR. These antibodies also bound to the LR, but only just anterior to its being joined by the “x” MT from the RR (Figures 3A, S5). The impression is made of the SFA signal “transferring” from MT “x” to the SR+LR combination. This suggests that SFA takes on an independent structural role here, in which it reinforces this junction. Our antibodies also stained three structures that were distinct from MTs: the B-fiber, the C-fiber, and the composite fiber. The B-fiber of *P. pyriformis* has a periodicity of 33 nm^23^. The composite fiber was not measured in that study, but the composite fiber of *P. eleionoma* has a 30-nm periodicity^24^. The C-fiber’s composition is less clear, as it was reported as unstriated in *P. pyriformis*^23^, but of 30-nm striation in *P. convexa*^29^, which is probably the same organism as *P. pyriformis*^24^. The C-fiber’s identity in our preparation is also uncertain: Although it has always been localized to the dorsal or “inner” side of the LR^3,23,29^, our results demonstrate that the distal end of the C-fiber often protrudes from the ventral or “outer” side of the LR (Figures 3A, 4A, 4B; Videos S1, S3). We feel the most probable interpretation of this is the C-fiber became dissociated with the LR, and either the distal end of the C-fiber was displaced ventrally or (more likely) the LR was displaced dorsally. Consistently with this interpretation, the C-fiber only projected ventrally in some cells (and usually in more-disrupted cells), and was parallel to the LR in others (Video S2). All these descriptions are compatible with these fibers being made of P8 or a combination of P7 and P8 proteins which self-assembled *in vitro* into filament bundles with 34.6 ± 2.2 nm periodicity of striation.

The B-fiber, C-fiber, and composite fiber are thought to be characteristic of “typical excavates”^3^, and homologs of these structures are not currently known in other organisms^3,5^. The B-fiber in particular appears to be a well-conserved structure, being attached only at its edges to the RR and consistently reported as having a 30-nm period of striation (reviewed by Simpson^3^). This implies that the B-fiber in other “typical excavates” comprises SFA as well. If this is the case, the argument for its homology across typical excavates will be greatly strengthened.

But that homology may extend beyond excavates. The narrower and wider roots of chlorophyte algae are thought to be homologous with the excavate LR (= R1) and RR (= R2), respectively^4,6^. System I fibers, which are built out of SFA^8^ and which were first described in green algae^30^, have a geometric relationship to these roots that is similar to that of the B-fiber to the RR and the C-fiber to the LR in excavates. In addition, the structure associated with the narrower root in chlorophytes is a multi-layered structure, as is the C-fiber in some ‘typical excavates’ — including, incidentally, *P. pyriformis*^23^. In other words, it is possible that the B- and C-fibers are not unique to excavates after all, being both compositionally and structurally homologous in organisms with cell architectures and phylogenetic relationships far removed from those of ‘typical excavates’.

The precise functions of SFAs seem hard to generalize: they may associate with individual or grouped MTs, or they may form structures separate from MTs, and while they are generally known from interphase cells, in chlorophytes and apicomplexans they are involved with cell division^13,31^. In this regard, System I fiber homologs built out of SFAs show a versatility similar to MTs themselves. But any genetic nature of this versatility is unknown. *Paratrimastix pyriformis* has (at least) five SFA paralogs, in all three paralog families, which may have subfunctionalised to account for different functional or structural roles. We were only able to investigate two within-lineage Group 1 paralogs in this study, and while our in-vitro TEM images show differences in the polymerization behavior of each, our antibodies were not specific enough to localize each differentially.

Still, we may propose some moderately confident hypotheses. Of the two paralogs we investigated, only P8 formed fibers *in vitro*, and those fibers had a ∼34-nm periodicity. It is natural to presume that it is P8 that contributes to striated structures with similar periodicities, namely the B-fiber, C-fiber and composite fiber. P7’s tendency to exist in a monomeric or oligomeric state might also be more consonant with its taking on the MT-binding role seen in our ExM images, given that no striations are known along the marked MTs. These hypotheses should be tested using more specific antibodies in the future.

Other preaxostylans also have two Group 1 paralogs. Our phylogenetic analysis suggests that each pair of paralogs arose independently in each preaxostylan species, or perhaps an ancestral duplication did not diversify until after individual lineages split off from the common preaxostylan stock. As such, there is no reason to suppose that any subfunctionalisation found in *P. pyriformis* will also be seen in any other species of this group. Other eukaryote lineages either have only a single Group 1 paralog or evolved their own paralogs independently. There is likewise no reason to assume that any allotment of functions to each paralog in preaxostylans is mirrored in any other group of eukaryotes. Cases in which the same division of SFA function is found in another group of eukaryotes with two Group 1 paralogs (the common ancestor of which had only one Group 1 paralog) are almost certainly convergent, and investigation of the biochemical properties in common between corresponding paralogs may be enlightening.

Again, we note that, while one or another paralog is unknown from several lineages within the “mega-group” Diaphoretickes, all “typical excavate” lineages possess SFA paralogs in Groups 1, 2a, and 2b (Figures S1, S2, S6). While we can only speculate at present, each of these deep-level paralogs may have shared functions across these lineages. There are several other fibrous structures in “typical excavate” cell architecture that remain uncharacterized. In particular, the I-fiber has a markedly periodic substructure^3^, and it may be that this structure contains Group 2 SFAs. To date, Group 2 paralogs have only been characterized in ciliates^9,11^, and their periods of striation are on the order of 30 nm^11,32^. This periodicity is similar to that of Group 1, so any functional differences between the two paralog families are probably independent of their striation patterns. All of this further emphasizes the localization of Group 2 paralogs in both *P. pyriformis* and other organisms as a prime goal for future studies.

While SFAs are known as structural proteins, we might propose nonstructural roles for them as well. In almost all flagellates, and in particular in all known “typical excavates,” cells do not divide symmetrically^33^. Rather, one daughter cell always receives the parental anterior flagellum (AF), and the other the parental posterior flagellum (PF), each flagellum with its associated structures. The daughter cell receiving the parental PF maintains that flagellum and associated roots as posterior structures, while in the other daughter cell, the AF transforms into a PF, and existing anterior roots transform into posterior roots. This means that PF-related structures are permanent, outlasting the independent lifetime of individual cells. Significantly, all of the structures tagged by our Group 1 SFA ABs are associated with the PF’s basal body. We might conjecture that the presence of SFAs on the iRR and iRMTs could provide a signal for stability, or could themselves physically stabilize the posterior-system MTs, to maintain them intact through the next division cycle. This, and other speculations on the roles of SFAs in maintaining cytoskeletal structures, must await further investigations.

## Conclusions and Future Directions

This study has explored the diversity of homologs of striated fiber assemblins (SFAs) across eukaryotes and determined the localization of two paralogs in *Paratrimastix pyriformis*. The localization was established using expansion microscopy, which, combined with specific immunostaining, provides insights into cellular structures at a combination of level of detail and context that is difficult to obtain through conventional fluorescence microscopy or electron microscopy alone. The data lead us to the conclusion that at least two paralogs of SFAs are present within the *P. pyriformis* feeding groove, where they decorate specific microtubules and build three striated fibers: the B-, C-, and composite fibers. The latter function is supported by the ability of one of these paralogs to form *in vitro* fibrous structures with striation similar to the mentioned fibers.

We suspect that SFAs also play a role in the development of cytoskeletal structures throughout the cell and flagellar cycles. SFA paralogs are present in all eukaryotes possessing “excavate” morphology, of which the groove is a central feature. It is thus reasonable to suspect that morphologically and positionally similar fibers in other organisms with such grooves, some of which are not closely related to *P. pyriformis* and probably branch on the opposite side from the root of the eukaryotic tree, will also be composed of SFAs. Should this be proven in the future, it would be a strong indication that SFAs, a protein family with broad distribution but diverse localizations in extant taxa, had an ancestral role in the feeding apparatus of the common ancestor of most or all eukaryotes. Overall, our work provides a glimpse into potential homologies of cytoskeletal structures across eukaryotes, setting the stage for further investigations into basal eukaryotic morphology and the architecture of the ancestral eukaryotic cell.

## MATERIALS AND METHODS

### Bioinformatics

*Data curation*. A “core” dataset was developed based on that of Lechtreck^13^, which contained only well-attested SFA sequences, specifically including those sequences experimentally confirmed to be SFAs (*Spermatozopsis similis* SFA, GenBank AAB26641.1; *Giardia intestinalis* β-giardin, GenBank P15518.2; *Toxoplasma gondii* SFA2, ToxoBase TGG_7332; *Phytophthora ramorum*, JGI-81943). To provide taxonomic breadth, the core dataset was expanded to include additional sequences, using data from Harper et al.^12^ and carefully selected sequences from GenBank (NCBI: https://www.ncbi.nlm.nih.gov/). To ensure the exclusion of spurious data, each core sequence was compared against all remaining core sequences using BLASTP^34^. All comparisons considered alignment length, percent identity, and E-value. Further amino-acid sequences were obtained from additional published datasets^9,11^, as well as data mined from the public databases GenBank and EukProt^35^. Taxonomically targetted data were mined from genome and transcriptome projects for the oxymonads *Monocercomonoides exilis*^36^, *Blattamonas nauphoetae*^37^, and *Streblomastix strix*^38^; the non-oxymonad preaxostylans *Paratrimastix pyriformis*^37^ and *Trimastix marina*^39^; and the fornicates *Retortamonas caviae* and *R. dobelli*^40^, *Ergobibamus cyprinoides, Aduncisulcus paluster, Dysnectes brevis*, and *Chilomastix cuspidata*^41^. All sequences were confirmed by reciprocal BLAST. When possible, partial sequences from one database were replaced with corresponding longer sequences from another, and only data from a single source was used for each sequence (i.e., partially overlapping sequences were not assembled into longer contigs). Sequences over ∼450 bases long were manually trimmed to within ∼150 residues of ‘core’ data, with methionine as the N-terminal residue when possible.

*Sequence alignment*. Initially, amino acid sequences were assembled into a single matrix and aligned using MUSCLE^42^, followed by manual alignment and masking. Preliminary trees were generated from masked alignments using RAxML v8.2.12^43^, using the LG substitution model and the CAT model of site variation, with support evaluated using 200–500 rapid bootstraps. Two primary clades of paralogs (Group 1 and Group 2) were identified early in this process (see Results); these paralogs were then divided into separate matrices and analyzed independently. Sequences that were not clearly recognizable in the alignment as belonging to a specific paralog were aligned individually to each paralog and assessed using rapid-bootstrap phylogenies as described above.

Long-branching sequences (defined as having branch lengths with ≥ 1.5 substitutions per site and being ≥ 2× the length of the next-longest sequence), near-identical sequences, and members of overrepresented clades (predominantly ciliates) were pruned from both matrices. The final Group 1 and Group 2 matrices had 158 and 144 sequences and 247 and 242 sites, respectively. Additionally, several combined matrices were assembled from Group 1 and Group 2 matrices, with the internal alignment of each paralog unchanged but the alignment between the clades varying by treatment.

These were evaluated using RAxML as described above; the combined dataset selected for further examination contained 302 sequences and 262 sites.

*Phylogeny*. Trees were generated from all three data sets using RAxML under the LG model with a gamma distribution of site rate variation and empirically determined base frequencies. The final topology in each case resulted from the best of 100 parallel runs, and support values were taken from 1,000 bootstrap replicates using the same model. Analyses were repeated using IQ-TREE^44^, with 100 parallel runs under the LG+F+G model to provide guide trees, and the LG+PMSF+C20+G model for final analyses, which included 200 bootstrap replicates. Each of the three final datasets was also subjected to Bayesian analysis using MrBayes v3.2.7^45^ to implement four parallel runs, each with eight chains, a heating parameter of 0.05 (Group 1) or 0.10 (Group 2 and combined data), two swaps every two generations, and a 10% burnin. These were repeated using the same parameters (with exceptions noted below) with the RAxML tree as a starting tree, with five perturbations of the tree applied to each chain, and a heating parameter of 0.10. The best tree from each of these analyses was used to seed a third Bayesian analysis, with three (Group 1) or two (Group 2 and combined data) perturbations per tree, four swaps per cycle (Group 1) or two swaps every two generations (Group 2 and combined data), and heating parameters of 0.05 (Group 1) or 0.10 (Group 2 and combined data). Since no runs had converged after 10^7^ generations, the run with the highest terminal likelihood was used for each analysis. As with RAxML analyses, Bayesian analyses used the LG substitution model, with a gamma distribution of site variation and empirically determined base frequencies.

*Structural analysis*. Protein-domain prediction was carried out on all sequences using the Simple Modular Architecture Research Tool (“SMART”) online platform^46^. Results from these analyses were processed through an in-house pipeline that (a) extracted and compiled predictions of coiled-coil and low-complexity regions as well as Pfam predictions for “SF-assemblin” and “Baculo_PEP_C” (the second-most-common Pfam prediction), (b) aligned, trimmed and masked per-site prediction results according to the Group 1 or Group 2 alignments, as appropriate, and (c), for easier visualization, mapped all predictions onto a single line (priority being given, in order, to coiled-coil domains, low-complexity domains, “SF-assemblin” predictions, and “Baculo_PEP_C” predictions) and collapsed lines by a factor of 7 (the number of residues required for one complete coil in a coiled-coil structure). Other reduction factors were also used. The factor-7 visualizations were aligned with their corresponding taxa in phylogenetic trees for visual comparison.

### Preparation of Recombinant Proteins

The recombinant proteins for antiserum production were prepared and purified as described in Peña-Diaz et al.^47^ Initially, the gene models for locus tags *PaPyr_7259* (P7) and *PaPyr_8006* (P8) were downloaded from the genome assembly^37^ and manually curated by mapping of transcripts. This curation resulted in the shortening of both gene models and their final coding sequences and translations are provided in Supplementary Document 1. The complete coding regions were amplified from *Paratrimastix pyriformis* cDNA using specific primers (PaPyr7259-F ATCGCATATGAGCACCCCGCCGCCCAGCTCTCCTTCCCC; PaPyr7259-RP ATATCTCGAGATGGGTGACCAGGTGCAGGCCCTCCTGCAG; PaPyr8006-F ATCGCATATGCAAGCCGAGACTCCGATTCCCACTC; PaPyr8006-R GCTAGCGGCCGCATGGGTGATGACGTGGAGGCCGTCCTG). Amplicons were cloned into the bacterial expression vector pET30a using the vectors’ C-terminal His-tags. Proteins were expressed in *E. coli* Rosetta2 DE3 cells by autoinduction for no more than 18 hours. Bacterial lysates were prepared by resuspending the cell pellets in HNGB (50 mM HEPES-KOH, pH 8.0, 300 mM NaCl, 8 M urea, 10% (v/v) glycerol, 2 mM beta-mercaptoethanol) in the presence of a protease inhibitor cocktail (Roche, Basel, Switzerland). These mixtures were incubated on ice for 30 min, then loaded into a 35-ml standard pressure cell and broken on a G-M high-pressure omogenizer at 10,000 kPa on a high ratio in three rounds. Whole-cell lysates were ultracentrifuged at 100,000 × *g* for 1 hour at 4° C in a SW40Ti rotor on a Beckman ultracentrifuge. Supernatants (clear lysate) were loaded in 20 ml columns packed with 5 ml of HisPur Ni-NTA resin (Thermo-Fischer, Waltham, Massachusetts, USA) equilibrated with HNGB. Columns were further washed with the same buffer, and proteins were eluted using 250 mM imidazole buffered in HNGB. Sample eluates were verified by SDS-PAGE and western blotting.

### Electron Microscopy

Proteins P7 and P8 were purified after ectopic expression in *E. coli* as described above, then re-natured by four steps of dialysis (25 mM HEPES-KOH, pH 7.5, 5 mM MgCl, 10 mM arginine, 10%(v/v) glycerol) over the course of 48 hr to remove the urea. Freshly dialyzed samples were stored at 4°C overnight before further processing. Samples were adsorbed on glow-discharged carbon-coated 400 Mesh Copper grids (Electron Microscopy Sciences #cF400-CU), negatively stained with uranyl acetate (4% aqueous solution), air-dried, and analyzed using a JEM2100-Plus Transmission Electron Microscope (JEOL) operating at 200 keV. Representative images were collected with TemCam–XF416 camera (TVIPS). For cryo-EM analysis, 4 µL of sample was first applied to glow-discharged Quantifoil cryo-EM grids (R 2/1, 300 mesh), incubated for 10 sec, then automatically blotted and vitrified in liquid ethane using LEICA EM GP2 plunge freezer. Cryo-EM data were then acquired in a semi-automated mode using SerialEM version 4.1.17^48^ at a pixel size of 1.44 Å/px and low dose exposure settings (< 50 e^−^/Å^2^) using the same microscope and camera as above.

### Preparation and Purification of Antibodies

The extracted proteins were used to raise polyclonal antibodies in rats as previously described^49^. Cyanogen-bromide-activated Sepharose 4B (C9142, Sigma-Aldrich, St. Louis, USA) was used for affinity purification of anti-P7 and -P8 antisera. Beads were re-swelled following the manufacturer’s instructions in ice-cold 1 mM HCl and washed with coupling buffer (0.1 M carbonate buffer, 0.5 M NaCl, pH 8.3). Purified *P. pyriformis* proteins were dialyzed against coupling buffer and incubated with the beads overnight at 4° C. Beads were collected and washed with coupling buffer, and subsequently incubated in 0.1 M Tris-HCl, pH 8.0, for 2 hr at room temperature. Beads were then washed for three cycles with Tris buffer (0.1 M Tris-HCl pH 8.0, 0.5 M NaCl) followed by acetate buffer (0.1 M sodium acetate, 0.5 M NaCl, pH 4.0) at 4° C. The antisera were then incubated with beads overnight at 4° C, and the bound antibodies were eluted by 0.2 M glycine, pH 2.8.

### Co-immunoprecipitation

Prior to immunoprecipitation, 2 l of well-grown *P. pyriformis* culture was filtered as previously described^50^. Briefly, a *P. pyriformis* culture in the exponential phase of growth was filtered to remove co-cultured bacteria, first through filter paper and then through a 3-µm polycarbonate membrane filter. Membrane filters were then washed twice with 10% (v/v) LB medium, and cells were collected by centrifugation at 1,000 × *g* for 10 min at 4° C. The resulting pellet was confirmed to contain 1*10^7^ *P. pyriformis* cells.

Immunoprecipitation was carried out using the Dynabeads Protein G Immunoprecipitation Kit (Thermo-Fisher, Waltham, Massachusetts, USA) according to the manufacturer’s instructions. Briefly, 1 ml of lysis buffer (25 mM Tris-HCl pH 7.5, 150 mM NaCl, 5 mM MgCl, and 0.1% (v/v) Triton X-100), supplemented with cOmplete EDTA-free protease inhibitors (Roche, Basel, Switzerland), was added to the pellet of *P. pyriformis* cells. This was incubated with rotation for 1 hr at 4 °C. Cleared lysate was centrifuged at 1,000 × *g* for 5 min at 4 °C, and the supernatant was transferred to a new 1.5-ml tube and kept on ice. The magnetic-beads/antibody complex was incubated with *P. pyriformis* cell lysate for 30 min with rotation at room temperature. For further western blot analysis, the pulled-down proteins were eluted under denaturation conditions. This procedure was repeated using RIPA lysis buffer (25 mM Tris-HCl pH 7.5, 150 mM NaCl, 1% (v/v) Triton X-100, 0.1% (v/v) SDS, and 0.5% (v/v) sodium deoxycholate) supplemented with cOmplete EDTA-free protease inhibitor (Roche).

### Expansion Microscopy

Cells were prepared according to the methods reported by Gorilak et al.^51^, with minor modifications.

#### Fixation

Cells were cultured and filtered as described above. Roughly 5 × 10^5^ cells per specimen were resuspended in sucrose buffer (25 mM sucrose; 10 mM HEPES-KOH, pH 7.5; 2 mM MgSO_4_) containing 4% formaldehyde and 4% acrylamide. Fixed cells were transferred directly to poly-L-lysine-coated coverslips, incubated overnight at 4° C, and then washed once with phosphate-buffered saline solution (PBS).

#### Gelation

Immediately prior to gelation, 0.5% N, N, N’, N’-tetramethylenediamine and ammonium persulphate were mixed with a monomer solution containing 19% sodium acrylate, 10% acrylamide, and 0.1% N, N’-methylenebisacrylamide in PBS. This was then dispensed in 50-µl aliquots onto glass coverslips in a wet chamber, on which coverslips containing cells were placed, cell side down. These were incubated on ice for 5 min and then at 37° C for 30 min.

#### Denaturation

Specimens were incubated at room temperature for a few minutes in denaturation buffer (50 mM Tris-HCl pH 9.0, 200 mM NaCl, 200 mM sodium dodecyl sulphate) until coverslips detached from gels. Gels were then transferred to 1.5-µl centrifuge tubes with fresh denaturation buffer and incubated for 1 hr at 95° C. After this, specimens were washed at room temperature with PBS thrice for 20 min each.

#### Antibody staining and expansion

Specimens were blocked with 2% bovine serum albumin in PBS for ≥ 1 hr at 4° C. For primary antibody staining, rat anti-P7 and anti-P8 sera, and guinea pig antitubulin (AA345, ABCD antibodies, Geneva, Switzerland), were diluted in 250 µl of blocking buffer at 1:50 and 1:100 titrations, respectively, and incubated overnight at 4° C. Gels were then washed thrice with PBS for 20 min at 4° C. Secondary antibodies (A11007, Alexa Fluor™ 594, goat anti-rat IgG; A11073, Alexa Fluor™ 488, goat anti-guinea pig IgG; Invitrogen, Waltham, Massachusetts, USA) were diluted 1:250 in blocking buffer, in which specimens were incubated overnight in the dark at 4° C. Afterward, specimens were washed thrice with PBS for 20 min at 4° C. Prepared gels were expanded by incubation in three changes of ultrapure water for 20 min each at 4° C.

#### Imaging and image processing

Expanded gels were transferred to poly-L-lysine-coated glass-bottomed dishes. Specimens were imaged using a Nikon CSU-W1 spinning disk field scanning confocal microscope (Nikon Instruments Inc., New York, USA) with a 63× water immersion objective. Voxel sizes were calculated to be 56 × 56 × 150 nm. Images were processed using Image-J 1.53k^52^.

## Supporting information

Supplementary file captions; supplementary figures S1-S6; sequence data used in paper

## RESOURCE AVAILABILITY

The data supporting all figures can be found in the published article and its online supplemental material. Alignments are available upon request.

## ACKNOWLEDGEMENTS

This work was supported by the Czech Science Foundation project number 23-07277S to VH and by Cooperatio Biology to VH. The authors would like to acknowledge the Imaging Methods Core Facility at BIOCEV, which is supported by the Ministry of Education, Youth and Sports of the Czech Republic (document number LM2023050 Czech-BioImaging), for their assistance in this study. Special thanks to Alastair GB Simpson for consulting the structure of the *P. pyriformis* cytoskeleton.

## AUTHOR CONTRIBUTIONS

Conceptualization, V. H. and A. A. H.; Methodology, A. A. H., Y-K. F. P. P-D. M. Z., J. M.; Investigation, A. A. H., A. K. J. Z., J. N., A. Š., E. Š., D. L, V. H.; Visualisation, A. A. H., A. K.; Writing – Original Draft A. A. H.; Writing – Review & Editing, A. A. H., P. P-D., D. L., V. H.; Funding Acquisition, V. H.; Supervision, V. H., P. P-D., D. L., M. Z.

