## Supplementary file captions; supplementary figures S1-S6; sequence data used in paper for "Striated fiber assemblins build the feeding groove of an excavate flagellate"

#### SUPPLEMENTAL INFORMATION TITLES AND LEGENDS

**Figure S1: Phylogeny of combined dataset containing all paralogs of striated fiber assembling, related to Figure 1.** Tree was generated from maximum-likelihood search using GTR+Γ model as implemented in RAxML; topology is from best of 100 starting random-taxon-addition parsimony trees. Support values at nodes are from 1000 nonparametric bootstrap replicates (above) and 200 bootstraps using LG+PMSF+C20+G model with 100 starting trees per replicate (below). Values < 50% are indicated by a dash ('–'), or omitted when both analyses returned < 50%. Black circles in branches indicate full support from Bayesian posterior probability; all branches without circles had probability scores < 0.75.

**Figure S2: Phylogenetic tree of Group 2 paralogs of striated fiber assembling, related to Figure 1.** Tree was generated from maximum-likelihood search using GTR+Γ model as implemented in RAxML; topology is from best of 100 starting random-taxon-addition parsimony trees. Support values along branches are from 1000 nonparametric bootstrap replicates (above branches) and from 200 bootstrap replicates using LG+PMSF+C20+G model with 100 starting trees per replicate (below). Values < 50% are indicated by a dash ('–'), or omitted when both analyses returned < 50%. Black circles in branches indicate full support from Bayesian posterior probability; open circles indicate support ≥ 0.95; no unlabelled branch was supported by Bayesian analysis (probability < 0.95). Upper (salmon) region represents Group 2a paralogs; lower (purple) region represents Group 2b. Ciliates, the only lineage in which Group 2 paralogs have been characterised, are highlighted in yellow. Numbers in gold shading indicate 'structural groups' defined by Nabi et al. 2019. Colored sequence names indicate previously characterised proteins. Figures at right are mappings of predicted coiled-coil domains onto sequence, with each character indicating seven amino acid sites (the number of sites generating a single coil); '@' indicates location of probable coiled-coil domains, '-' indicates sequence with no coiled-coil domain prediction; absence of symbol within mapping indicates gap in alignment (i.e., deletion); absence of symbol at ends of mapping indicates sequence not available.

**Figure S3. Ultrastructural analysis of P7, P8, and P7+P8 protein samples by cryo-electron microscopy, related to Figure 2. A:** Homogeneous distribution of P7 (likely dimers or oligomers) with globular (head) and stretched (tail) domains discernible in some projections. **B:** Protofilaments, bundles and lateral assemblies formed by P8. **C–D:** Mixture of P7 and P8 assembled either into two-dimensional sheets of laterally-aligned filaments (C), or into three-dimensional structures composed of equilateral triangles (D). Note that length of striation units (in C) is close or equal to triangle side length (in D) (approximately 40 nm). All images are pairs of overview images (on left) with enlarged details (on right). Scale bar = 50 µm.

**Figure S4: Western blot of antibodies raised against striated fiber assemblin proteins from *Paratrimastix pyriformis*, related to Figures 2 and 3. Left:** P7, **right:** P8. Arrows indicate location of SFA proteins.

**Figure S5: Expansion microcopy of whole cell of *Paratrimastix pyriformis*, related to Figure 3.** Three cells viewed from ventral-right side, stained with antibodies to structural proteins. **Top:** striated fiber assemblin paralog *PaPyr7259*; **middle:** tubulin; **bottom:** merged images. Note that some flagella have broken off close to their insertion points. Note also that singlet root is not visible as independent structure. Key to annotations: 'AF' = anterior flagellum; 'AR' = anterior root; 'B' = B-fiber; 'C' = C-fiber; 'CF' = composite fiber; 'CR' = compound root; 'fan' = dorsal fan; 'iRMT' = inner right root microtubules; 'iRR' = inner right root; 'LF' = left flagellum; 'PF' = posterior flagellum; 'oRR' = outer right root; 'RF' = right flagellum; 'SR+LR+C' = compound root comprising singlet root, left root and C fibre; 'x' = 'x' microtubule in iRMT. Scale bar ~ 5 µm (approximated based upon the diameter of flagellar axonemes and size of cell).

**Figure S6: Summary trees showing known distribution of paralogs of striated fiber assemblin in major lineages of eukaryotes.** Black lines indicate lineages known to possess at least one copy of paralog represented by tree; grey lines indicate no known paralogs in lineage. Light-colored background indicates eukaryotic 'megagroups' that may be paraphyletic; dark-colored arcs at edges of tree indicate eukaryotic 'supergroups' that are expected to be holophyletic.

**Table S1: Results of co-immunoprecipitation with P7 antibody, related to Figure S4.** List of proteins identified after co-immunoprecipitation of sample with fold change relative to negative control (beads without antibody). Dash ('–') indicates cases where protein was not identified in negative control. Striated fiber assemblins are shown in red.

**Table S2: Results of co-immunoprecipitation with P8 antibody, related to Figure 4.** List of proteins identified after co-immunoprecipitation of sample with fold change relative to negative control (beads without antibody). Dash (‘–’) indicates cases where protein was not identified in negative control. Striated fiber assemblins are shown in red.

**Table S3: Results of co-immunoprecipitation with P7 and P8 antibodies in RIPA buffer, related to Figures 4 and S4.** List of proteins identified after co-immunoprecipitation of sample with fold change relative to negative control (beads without antibody). Dash (‘–’) indicates cases where protein was not identified in negative control. Striated fiber assemblins are shown in red.

**Video S1: Animation of expanded cell of *Paratrimastix pyriformis* tagged with antibodies to tubulin (green) and striated fiber assemblin (magenta), related to Figure 4.** Cell is oriented to rotate about its anteroposterior axis, with anterior to top of screen. Signal from antitubulin antibody shown in green, signal from anti-P8 antibody in magenta. Animation is of the same cell as shown in Figure 4.

**Video S2: Animation of expanded cell of *Paratrimastix pyriformis* tagged with antibodies to tubulin (green) and striated fiber assemblin (magenta), related to Figure S4** Cell is oriented to rotate about its anteroposterior axis, with anterior to top of screen. Signal from antitubulin antibody shown in green, signal from anti-P7 antibody in magenta. Animation is of same cell as shown in Figure S4.

**Video S3: Video composed of all optical slices from the cell visualized in Figure 5, related to Figure 5.** Cell is viewed in cross-section, with dorsal to top of page and cell’s left to viewer’s right. Signal from antitubulin antibody shown in green, signal from anti-P8 antibody in magenta.

**Figure S1**

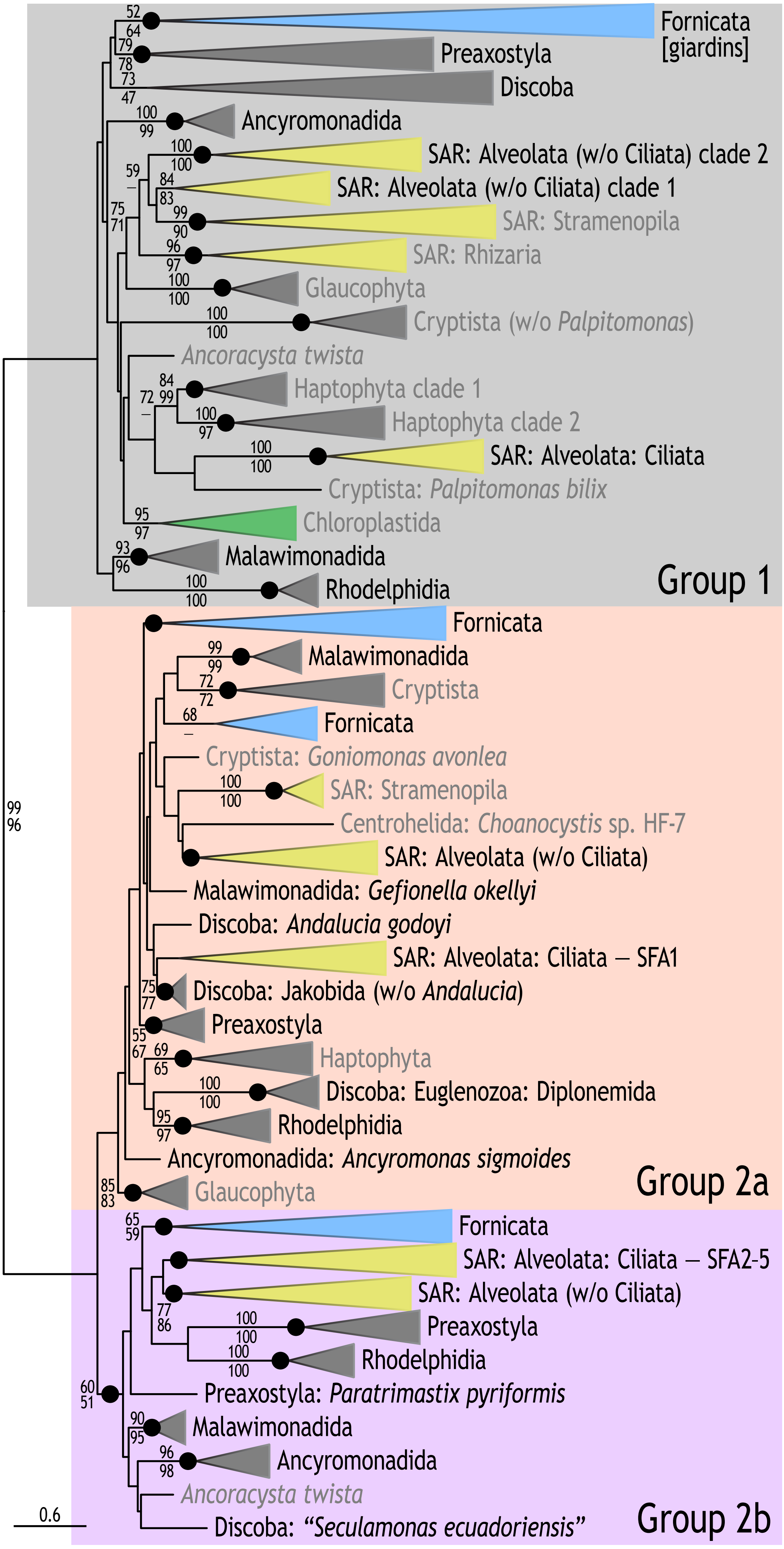
Fornicata [giardins]
Preaxostyla
Discoba
Ancyromonadida
SAR: Alveolata (w/o Ciliata) clade 2
SAR: Alveolata (w/o Ciliata) clade 1
SAR: Stramenopila
SAR: Rhizaria
Glaucophyta
Cryptista (w/o Palpitomonas)
Ancoracysta twista
Haptophyta clade 1
Haptophyta clade 2
SAR: Alveolata: Ciliata
Cryptista: Palpitomonas bilix
Chloroplastida
Malawimonadida
Rhodelphidia
Group 1
Fornicata
Malawimonadida
Cryptista
Fornicata
Cryptista: Goniomonas avonlea
SAR: Stramenopila
Centrohelida: Choanocystis sp. HF-7
SAR: Alveolata (w/o Ciliata)
Malawimonadida: Gefionella okellyi
Discoba: Andalucia godoyi
SAR: Alveolata: Ciliata – SFA1
Discoba: Jakobida (w/o Andalucia)
Preaxostyla
Haptophyta
Discoba: Euglenozoa: Diplonemida
Rhodelphidia
Ancyromonadida: Ancyromonas sigmoides
Glaucophyta
Group 2a
Fornicata
SAR: Alveolata: Ciliata – SFA2-5
SAR: Alveolata (w/o Ciliata)
Preaxostyla
Rhodelphidia
Preaxostyla: Paratrimastix pyriformis
Malawimonadida
Ancyromonadida
Ancoracysta twista
Discoba: "Seculamonas ecuadoriensis"
Group 2b
0.6

**Figure S2**

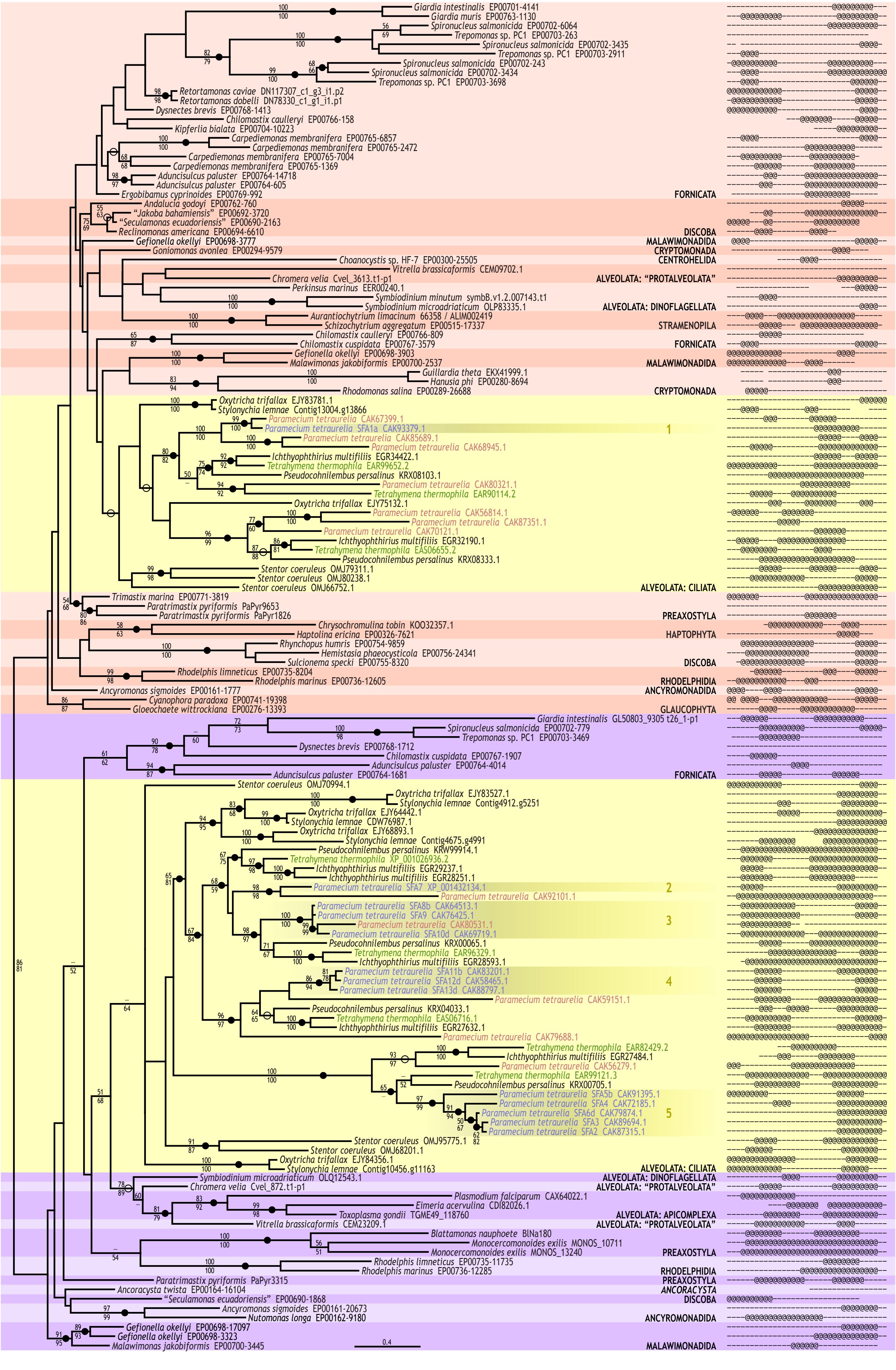

Giardia intestinalis EP00701-4141
Giardia muris EP00763-1130
Spironucleus salmonicida EP00702-6064
Trepomonas sp. PC1 EP00703-263
Spironucleus salmonicida EP00702-3435
Trepomonas sp. PC1 EP00703-2911
Spironucleus salmonicida EP00702-243
Spironucleus salmonicida EP00702-3434
Trepomonas sp. PC1 EP00703-3698
Retortamonas caviae DN117307_c1_g3_i1.p2
Retortamonas dobelli DN78330_c1_g1_i1.p1
Dysnectes brevis EP00768-1413
Chilomastix caulleryi EP00766-158
Kipferlia bialata EP00704-10223
Carpediemonas membranifera EP00765-6857
Carpediemonas membranifera EP00765-2472
Carpediemonas membranifera EP00765-7004
Carpediemonas membranifera EP00765-1369
Aduncisulcus paluster EP00764-14718
Aduncisulcus paluster EP00764-605
Ergobibamus cyprinoides EP00769-992
FORNICATA
Andalucia godoyi EP00762-760
"Jakoba bahamiensis" EP00692-3720
"Seculamonas ecuadoriensis" EP00690-2163
Reclinomonas americana EP00694-6610
DISCOBA
Gefionella okellyi EP00698-3777
MALAWIMONADIDA
Goniomonas avonlea EP00294-9579
CRYPTOMONADA
Choanocystis sp. HF-7 EP00300-25505
CENTROHELIDA
Vitrella brassicaformis CEM09702.1
Chromera velia Cvel_3613.t1-p1
ALVEOLATA: "PROTALVEOLATA"
Perkinsus marinus EER00240.1
Symbiodinium minutum symbB.v1.2.007143.t1
Symbiodinium microadriaticum OLP83335.1
ALVEOLATA: DINOFLAGELLATA
Aurantiochytrium limacinum 66358 / ALIM002419
Schizochytrium aggregatum EP00515-17337
STRAMENOPILA
Chilomastix caulleryi EP00766-809
Chilomastix cuspidata EP00767-3579
FORNICATA
Gefionella okellyi EP00698-3903
Malawimonas jakobiformis EP00700-2537
MALAWIMONADIDA
Guillardia theta EKX41999.1
Hanusia phi EP00280-8694
Rhodomonas salina EP00289-26688
CRYPTOMONADA
Oxytricha trifallax EJY83781.1
Stylonychia lemnae Contig13004.g13866
Paramecium tetraurelia CAK67399.1
Paramecium tetraurelia SFA1a CAK93379.1
1
Paramecium tetraurelia CAK85689.1
Paramecium tetraurelia CAK68945.1
Ichthyophthirius multifiliis EGR34422.1
Tetrahymena thermophila EAR99652.2
Pseudocohnilembus persalinus KRX08103.1
Paramecium tetraurelia CAK80321.1
Tetrahymena thermophila EAR90114.2
Oxytricha trifallax EJY75132.1
Paramecium tetraurelia CAK56814.1
Paramecium tetraurelia CAK87351.1
Paramecium tetraurelia CAK70121.1
Ichthyophthirius multifiliis EGR32190.1
Tetrahymena thermophila EAS06655.2
Pseudocohnilembus persalinus KRX08333.1
Stentor coeruleus OMJ79311.1
Stentor coeruleus OMJ80238.1
Stentor coeruleus OMJ66752.1
ALVEOLATA: CILIATA
Trimastix marina EP00771-3819
Paratrimastix pyriformis PaPyr9653
Paratrimastix pyriformis PaPyr1826
PREAXOSTYLA
Chrysochromulina tobin KOO32357.1
Haptolina ericina EP00326-7621
HAPTOPHYTA
Rhynchopus humris EP00754-9859
Hemistasia phaeocysticola EP00756-24341
Sulcionema specki EP00755-8320
DISCOBA
Rhodelphis limneticus EP00735-8204
Rhodelphis marinus EP00736-12605
RHODELPHIDIA
Ancyromonas sigmoides EP00161-1777
ANCYROMONADIDA
Cyanophora paradoxa EP00741-19398
Gloeochaete wittrockiana EP00276-13393
GLAUCOPHYTA
Giardia intestinalis GL50803_9305 t26_1-p1
Spironucleus salmonicida EP00702-779
Trepomonas sp. PC1 EP00703-3469
Dysnectes brevis EP00768-1712
Chilomastix cuspidata EP00767-1907
Aduncisulcus paluster EP00764-4014
Aduncisulcus paluster EP00764-1681
FORNICATA
Stentor coeruleus OMJ70994.1
Oxytricha trifallax EJY83527.1
Stylonychia lemnae Contig4912.g5251
Oxytricha trifallax EJY64442.1
Stylonychia lemnae CDW76987.1
Oxytricha trifallax EJY68893.1
Stylonychia lemnae Contig4675.g4991
Pseudocohnilembus persalinus KRW99914.1
Tetrahymena thermophila XP_001026936.2
Ichthyophthirius multifiliis EGR29237.1
Ichthyophthirius multifiliis EGR28251.1
Paramecium tetraurelia SFA7 XP_001432134.1
2
Paramecium tetraurelia CAK92101.1
Paramecium tetraurelia SFA8b CAK64513.1
Paramecium tetraurelia SFA9 CAK76425.1
Paramecium tetraurelia CAK80531.1
3
Paramecium tetraurelia SFA10d CAK69719.1
Pseudocohnilembus persalinus KRX00065.1
Tetrahymena thermophila EAR96329.1
Ichthyophthirius multifiliis EGR28593.1
Paramecium tetraurelia SFA11b CAK83201.1
Paramecium tetraurelia SFA12d CAK58465.1
4
Paramecium tetraurelia SFA13d CAK88797.1
Paramecium tetraurelia CAK59151.1
Pseudocohnilembus persalinus KRX04033.1
Tetrahymena thermophila EAS06716.1
Ichthyophthirius multifiliis EGR27632.1
Paramecium tetraurelia CAK79688.1
Tetrahymena thermophila EAR82429.2
Ichthyophthirius multifiliis EGR27484.1
Paramecium tetraurelia CAK56279.1
Tetrahymena thermophila EAR99121.3
Pseudocohnilembus persalinus KRX00705.1
Paramecium tetraurelia SFA5b CAK91395.1
Paramecium tetraurelia SFA4 CAK72185.1
Paramecium tetraurelia SFA6d CAK79874.1
5
Paramecium tetraurelia SFA3 CAK89694.1
Paramecium tetraurelia SFA2 CAK87315.1
Stentor coeruleus OMJ95775.1
Stentor coeruleus OMJ68201.1
Oxytricha trifallax EJY84356.1
Stylonychia lemnae Contig10456.g11163
ALVEOLATA: CILIATA
Symbiodinium microadriaticum OLQ12543.1
ALVEOLATA: DINOFLAGELLATA
Chromera velia Cvel_872.t1-p1
ALVEOLATA: "PROTALVEOLATA"
Plasmodium falciparum CAX64022.1
Eimeria acervulina CDI82026.1
Toxoplasma gondii TGME49_118760
ALVEOLATA: APICOMPLEXA
Vitrella brassicaformis CEM23209.1
ALVEOLATA: "PROTALVEOLATA"
Blattamonas nauphoete BlNa180
Monocercomonoides exilis MONOS_10711
Monocercomonoides exilis MONOS_13240
PREAXOSTYLA
Rhodelphis limneticus EP00735-11735
Rhodelphis marinus EP00736-12285
RHODELPHIDIA
Paratrimastix pyriformis PaPyr3315
PREAXOSTYLA
Ancoracysta twista EP00164-16104
ANCORACYSTA
"Seculamonas ecuadoriensis" EP00690-1868
DISCOBA
Ancyromonas sigmoides EP00161-20673
Nutomonas longa EP00162-9180
ANCYROMONADIDA
Gefionella okellyi EP00698-17097
Gefionella okellyi EP00698-3323
Malawimonas jakobiformis EP00700-3445
MALAWIMONADIDA
0.4

**Figure S3**

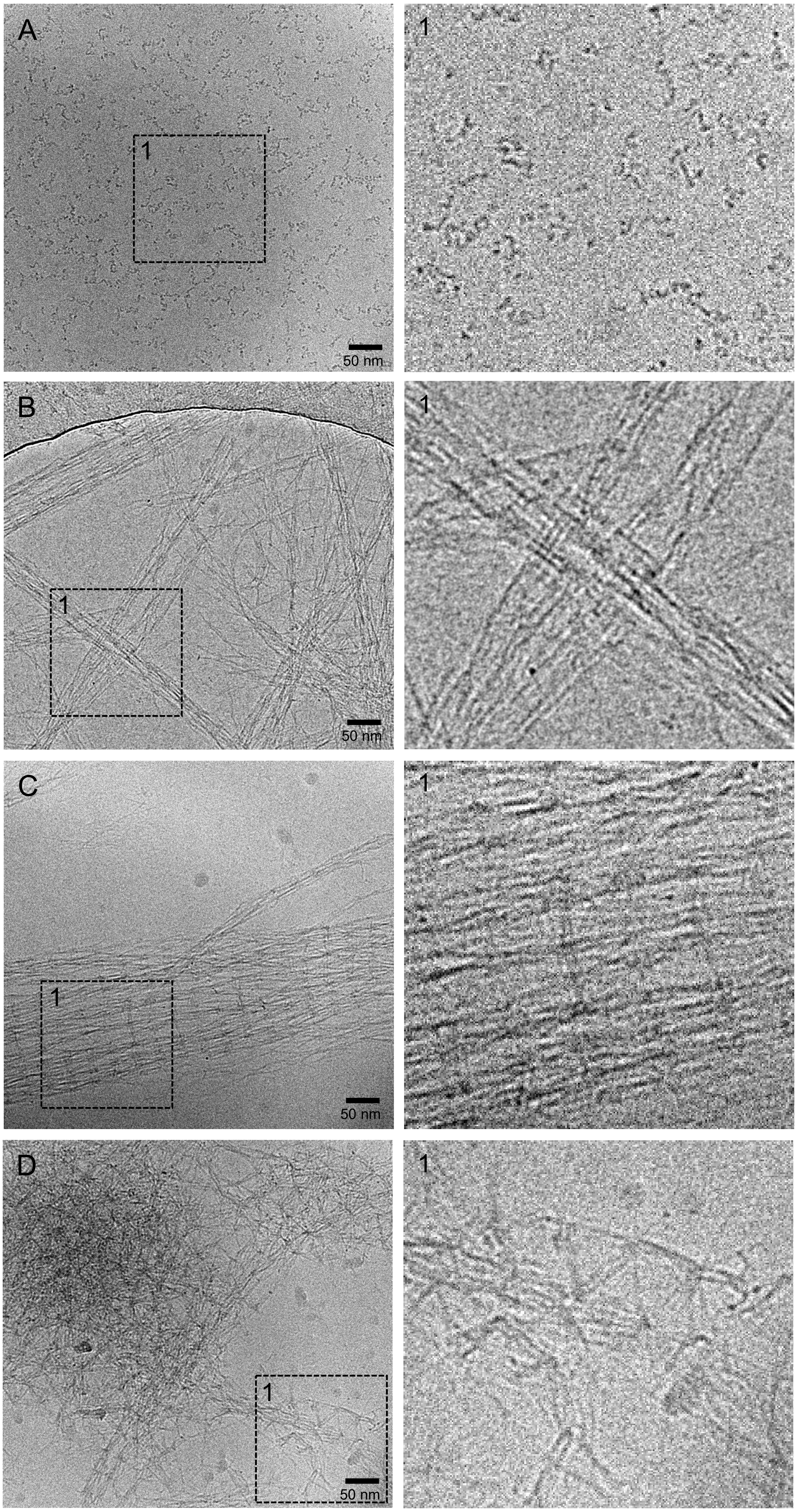

Figure S4

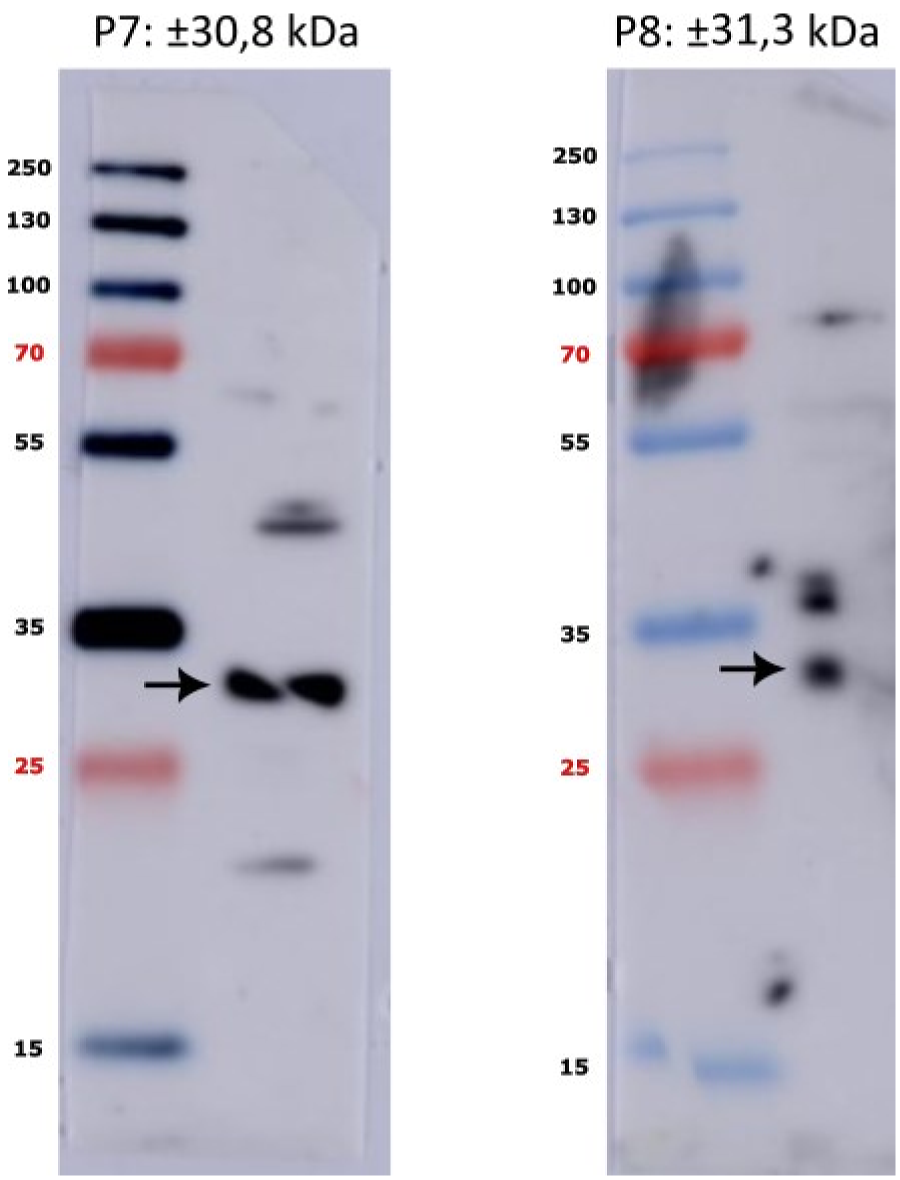

P7: ±30,8 kDa
250
130
100
70
55
35
25
15
P8: ±31,3 kDa
250
130
100
70
55
35
25
15

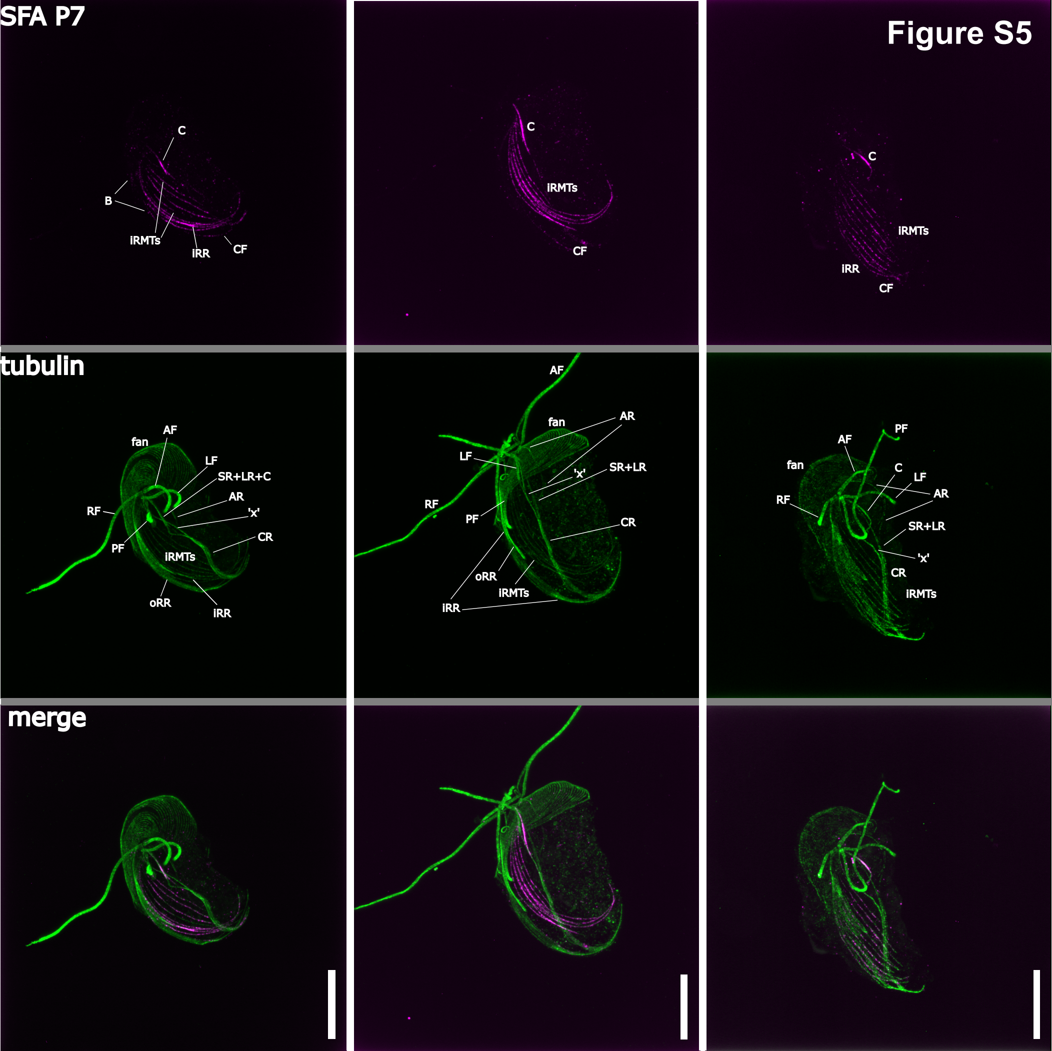
SFA P7
Figure S5
C
B
iRMTs
iRR
CF
C
iRMTs
CF
C
iRMTs
iRR
CF
tubulin
fan
AF
LF
SR+LR+C
AR
'x'
RF
PF
CR
iRMTs
oRR
iRR
AF
fan
AR
LF
'x'
SR+LR
RF
PF
CR
oRR
iRMTs
iRR
PF
AF
fan
C
LF
AR
RF
SR+LR
'x'
CR
iRMTs
merge

### Figure S6

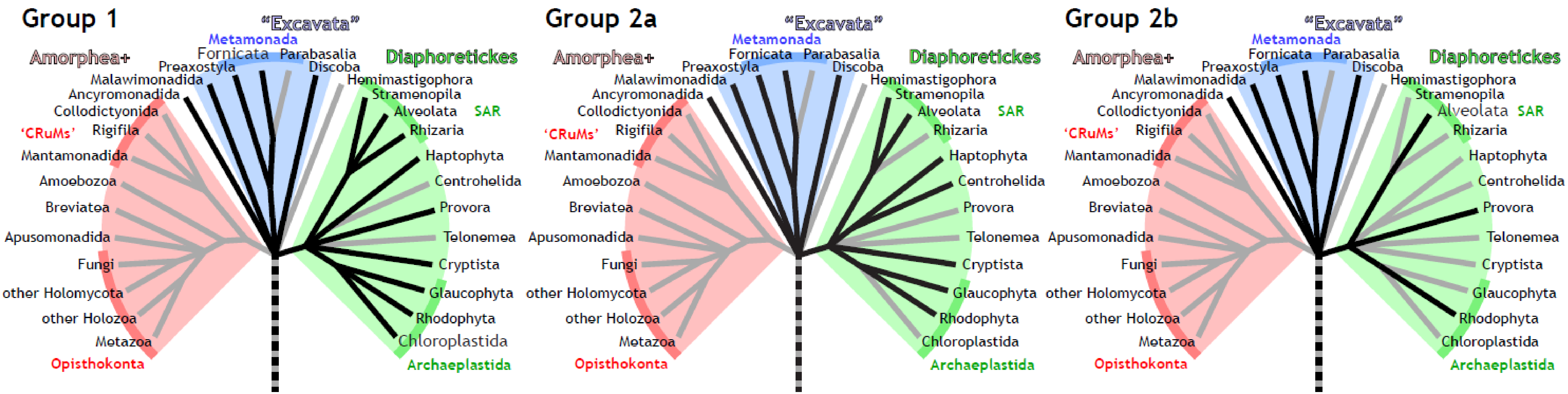
Group 1
"Excavata"
Metamonada
Fornicata
Parabasalia
Preaxostyla
Discoba
Amorphea+
Diaphoretickes
Malawimonadida
Hemimastigophora
Ancyromonadida
Stramenopila
Collodictyonida
Alveolata
SAR
'CRuMs'
Rigifila
Rhizaria
Mantamonadida
Haptophyta
Amoebozoa
Centrohelida
Breviatea
Provora
Apusomonadida
Telonemea
Fungi
Cryptista
other Holomycota
Glaucophyta
other Holozoa
Rhodophyta
Metazoa
Chloroplastida
Opisthokonta
Archaeplastida
Group 2a
Group 2b

**Supplementary document 1 – Corrected coding sequences of PaPyr7259 and PaPyr8006**

>PaPyr7259

ATGAGCACCCCGCCGCCCAGCTCTCCTTCCCCTCAGACTCCCCACAAGGACCAGTTGTCCCCCCT
CACCCAGCGGCTTTCCCTCTTGCAGAGCCGCTTTTCCAGTTTCCACTCTGATATCGAAACGGAGGCT
CAGCAACGTCGTGCGGACGACGAGATTCGTGTTCAGGCTCTTCGCGAAGTTGTTACCAAGATCGAGA
AGAACCTGACCCTGGAAATTAAAAGGCGGCAAGAAGCCGATCGCACTCTTCAGGGACTGATTGAACA
GCGAGTGACGGGATTGCAGGAAGCTCTCGAGACTGACCTGAAGGATAAGCTCCGCCAGATCATGAC
TGCCGTTGAATCCCTGAACACCCGGATCACCCAGCTCGAGCACGAGATTCGTGACGAGAAGGAGTG
CCGCCAAAAGGAGGTTGAGGACACCAACGCGATCCTTCTTCGCCAGCTGAACCAACTCCAGAACAC
GTTTGAGATGGAGAAGATGGCCCGGCTGGAACGCGAGAACCAGATTGTGAAGCGAGCCGGGGACG
ACTGCTTCCGTCTGCAGGAACGCATGGACGCCGAGAAGGTGGGCCGTGAGGCGGCTCTGACCCAG
CTCCGGGATGACTGGTCCGAGAACGAGCGCGTCCGGGGTAAGGCCGACGAGCAGTTCCGGGCGC
ATGTCCTCGGGGAGATAGCCGCACTGCACAACATCCTGCAGACCGAGCAGCAGAGCCGCGAGCAG
TCCGAGGCCCACATTGTGGCCACCGTCGACGAGATCGCGGCACGGCTGCAGGAGGGCCTGCACC
TGGTCACCCATTAG

>PaPyr7259

MSTPPPSSPSPQTPHKDQLSPLTQRLSLLQSRFSSFHSDIETEAQQRRADDEIRVQALREVVTKIEKNLTLE
IKRRQEADRTLQGLIEQRVTGLQEALETDLKDKLRQIMTAVESLNTRITQLEHEIRDEKECRQKEVEDTNAIL
LRQLNQLQNTFEMEKMARLERENQIVKRAGDDCFRLQERMDAEKVGREAALTQLRDDWSENERVRGK
ADEQFRAHVLGEIAALHNILQTEQQSREQSEAHIVATVDEIAARLQEGLHLVTH*

>PaPyr8006_1

ATGCAAGCCGAGACTCCGATTCCCACTCCCGCAACCCCGTCCTCTCGTTCTCAAGTCGATGGGACT
CCAGCTGCTTCCCCGACTGTCCAGCGCCTGCAGATGCTCTCCGAACGTTTCTCTGGGTTCCACACTG
ATCTGGAGACCGAGGCTCAACAGAGGCGACAGGACGATGAAATTCGTGTCCAGTCCATTCGTGATGTT
GTCTCCAAAATCGAAAAGAACTTGACGATGGAAATTAAAAGGAGGCAGGAAGCTGACCGTGCACTTCA
AAACCTTCTGGAAATGAGGGTGAATTCGCTGCAGGAAGCACTCGAATCCGATTTGAAGGATAAACTTC
GCCAGATTATGACGGCTGTTGAATCTCTCAATGGTAGGGTTCAACAAATCGAGCATGATGTCAAGGACG
AAAAAGAATCCCGGCAAAAGGAACTCGAAGACACGAATGCCCTCCTCCTTCGGCAACTGAACACTCT
GCAGAACACTTTCGAGATGGAGAAAATGTCCCGCATGGAGCGCGAGACGCAGATTCTCAAGCGGAT
GAGCGACGATGTCTTCCGGCTGCAAGAGAAAATCGACGGGGAGAAGCTGGCCCGCGAGACTGGG
CTTGCCCAGCTGCGGGATGAGATGCAGGAGGGTGCCCAGGCGCGCACCAAGGTCGACGAGAAGA
TGCGGGCCCACTTCCTGGAGGAGATCGCCTCCCTCCACGCCCAGTTGCAGGCTGAGCAGCAGGC
ACGCGAGGCCAACGAGGAACAGATTGTGGCGACTGTCGAGGATGTGGTATCGGGCCTGCAGGACG
GCCTCCACGTCATCACCCATTAA

>PaPyr8006_1

MQAETPIPTPATPSSRSQVDGTPAASPTVQRLQMLSERFSGFHTDLETEAQQRRQDDEIRVQSIRDVVSKI
EKNLTMEIKRRQEADRALQNLLEMRVNSLQEALESDLKDKLRQIMTAVESLNGRVQQIEHDVKDEKESR
QKELEDTNALLLRQLNTLQNTFEMEKMSRMERETQILKRMSDDVFRLQEKIDGEKLARETGLAQLRDEM
QEGAQARTKVDEKMRAHFLEEIASLHAQLQAEQQAREANEEQIVATVEDVVSGLQDGLHVITH*
